# Macrophages form integrin-mediated adhesion rings to pinch off surface-bound objects for phagocytosis

**DOI:** 10.1101/2023.08.01.551462

**Authors:** Kaushik Pal, Subhankar Kundu, Xuefeng Wang

## Abstract

Macrophages engulf micron-sized objects including pathogens and cell debris by phagocytosis, serving a fundamental role in immune defense and homeostasis^1, 2^. Although the internalization process of suspended particles has been thoroughly investigated^3, 4^, it is incompletely understood how macrophages internalize surface-bound objects by overcoming the surface binding. Here, we prepared a force-sensing platform which visualizes cell-substrate adhesive force by fluorescence. Macrophages are tested on this platform with micron-sized objects (E. coli, microbeads and silver nanorods) immobilized. By co-imaging integrin-transmitted forces and corresponding structural proteins, we discovered that macrophages consistently form integrin-mediated adhesion structures on the surface to encircle and pinch off surface-bound objects. We termed these structures phagocytic adhesion rings (PAR) and showed that integrin tensions in PARs are resulted from local actin polymerization, but not from myosin II. We further demonstrated that the intensity of integrin tensions in PARs is correlated with the object surface-bound strength, and the integrin ligand strength (dictating the upper limit of integrin tensions) determines the phagocytosis efficiency. Collectively, this study revealed a new phagocytosis mechanism that macrophages form PARs to provide physical anchorage for local F-actin polymerization that pushes and lifts off surface-bound objects during phagocytosis.

---

Through phagocytosis, phagocytes such as macrophages, neutrophils and dendritic cells engulf and digest micron-sized objects including bacteria, fungi, cell debris, mineral particles, etc. For suspended objects in solution, phagocytes deploy phagocytic receptors which bind and pull the objects to initiate phagocytosis^5-7^. Subsequently, phagocytes remodel plasma membrane and cytoskeleton to wrap on the objects and facilitate the internalization. In contrast to suspended objects, how surface-bound ones are internalized by phagocytosis is much less known. Surface-bound pathogens and materials are common in metazoan animal bodies. Many pathogenic bacteria adhere to the host before invading underlying tissues^8, 9^. Candida spp., the pathogenic fungi causing most frequent fungal infection in human, express adhesin^10^ to adhere tightly to human skin, and to endothelial and epithelial mucosal host tissues^11^. Cell debris, including neutrophil extracellular traps^12^, may stick to tissues. To internalize these surface-bound objects during phagocytosis, phagocytes need to overcome the adhesion of these objects on the surface. However, studies regarding the phagocytosis of surface-bound objects are scarce. Although one previous study showed that macrophages use filopodia to hook and shovel a remote E. coli for surface detachment^13^, the general mechanism remains unclear for the phagocytosis of surface-bound objects under the cell bodies of macrophages.

Because macrophages are adherent cells, we hypothesized that integrins, the major protein mediating macrophage adhesion, may play a role in the phagocytosis of surface-bound objects. Integrins have already been known to participate in the phagocytosis of suspended particles. For example, integrin α_M_β_2_, a subfamily of integrins highly expressed in macrophages, can function as a phagocytic receptor which binds to ligand-expressing pathogens or opsonized particles to facilitate their internalization^14, 15^. Integrins have also been shown to form a barrier to exclude phosphatase from phagocytic cups to facilitate phagocytosis maturation^16^. However, these studies were focused on the phagocytosis of suspended particles. The role of integrins remains unknown in the phagocytosis of surface-bound objects.

## Preparation of a force-sensing platform immobilized with micron-sized objects

In this work, we applied integrative tension sensor (ITS) to image integrin tensions produced by macrophages on a flat substrate during the phagocytosis of surface-bound objects. ITS is a tension sensor that visualizes integrin tensions by fluorescence at a sub-micron resolution^17-19^. The main construct of ITS is a nucleic acid duplex decorated with an integrin peptide ligand (Arg-Gly-Asp, or RGD)^20^, a fluorophore-quencher pair, and a biotin tag for surface immobilization (Fig. 1A). As integrins in adherent cells bind to the RGD ligand and transmit cell-generated force to ITS, the duplex can be mechanically unzipped and the fluorophore would be freed from being quenched, hence reporting the force by fluorescence. In this study, DNA/PNA (peptide nucleic acid)^21^ duplex instead of double-stranded DNA was adopted to construct ITS for its higher resistance to membrane DNase, which was expressed substantially on the cell membrane of macrophages^22^. ITS was grafted on glass surfaces at a surface density about 1000 μm^-2^ (Extended Data Fig. 1). The surfaces were also co-coated with fibronectin to support normal cell adhesion. *E*. coli, polystyrene beads and silver nanorods were immobilized on the surfaces to test the phagocytosis of surface-bound objects.

**Fig. 1.**
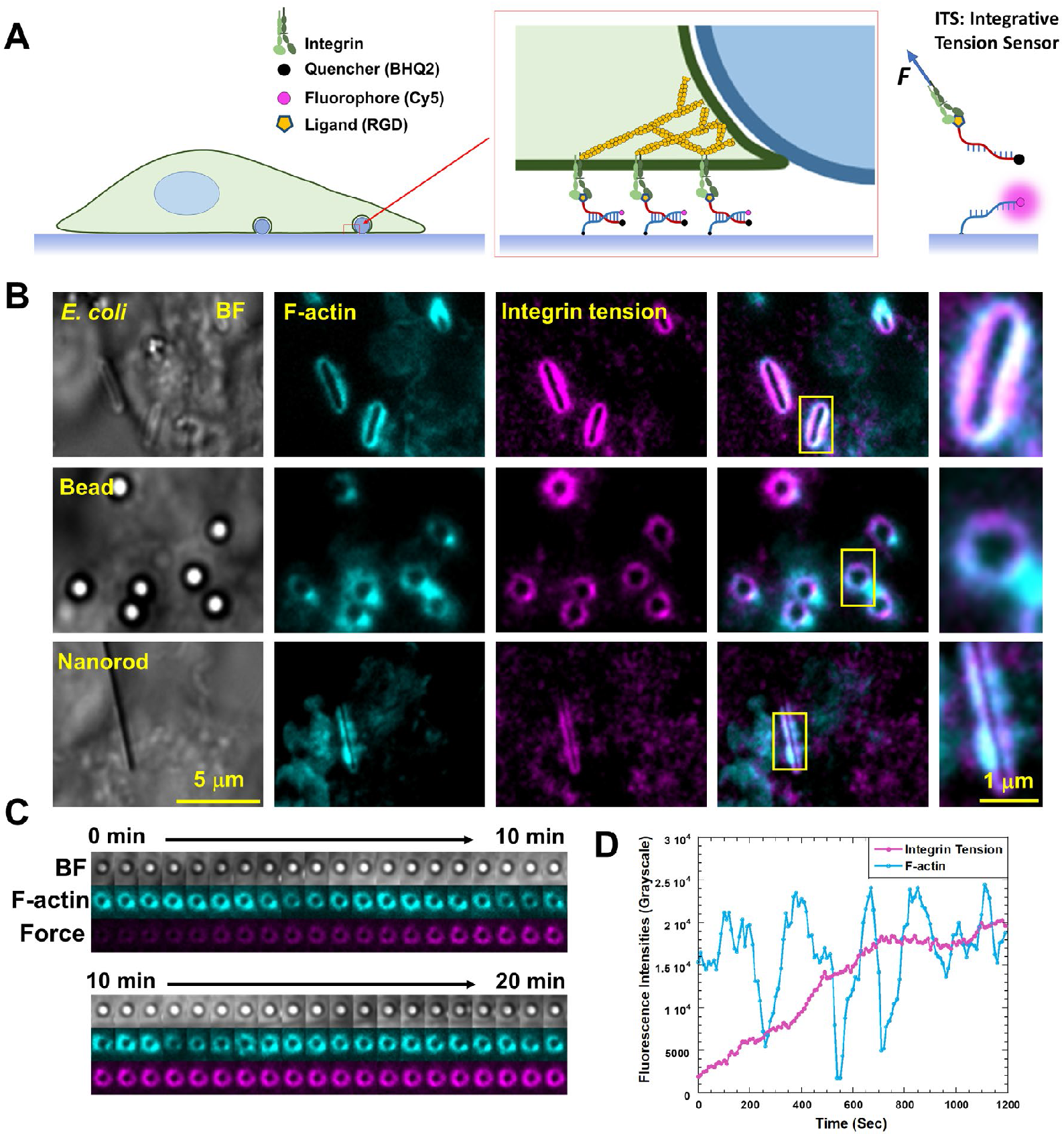
Macrophages form force-bearing adhesion structures surrounding surface-bound objects. (**A**) Integrin tensions in macrophages were imaged on a substrate grafted with integrative tension sensor (ITS) and immobilized with surface-bound objects. ITS is a DNA/PNA construct conjugated with an integrin ligand and a quencher-dye pair. ITS becomes fluorescent after integrin tension ruptures the duplex, hence recording integrin tension signal by fluorescence. (**B**) Macrophages induced from RAW267.4 cells consistently exhibited F-actin and integrin tension signal surrounding the surface-bound objects, including E. coli, micron-sized beads and silver nanorods. (**C**) Time-series co-imaging of F-actin and integrin tensions on a bead with a diameter of 1 μm (Supplementary Video 1). (**D**) F-actin exhibits periodic dynamics around the bead. Integrin tension signal is accumulated by time.

## Macrophages form force-bearing adhesion structures encircling surface-bound objects

On the force-sensing platform described above, mouse macrophages induced from RAW264.7 cells (Fig. 1B), human macrophages induced from THP-1 cells (Extended Data Fig. 2) and macrophages extracted from fish (Extended Data Fig. 3) were subsequently plated. The results show that these cells responded to the surface-bound objects and F-actin was recruited to surround the objects (Fig. 1B), suggesting the phagocytosis was initiated. Remarkably, cells also consistently and specifically produced integrin tensions encircling the objects regardless of the tested sizes and shapes of the objects (Fig. 1B, Extended Data Fig. 2 and Supplementary Video 1). These integrin tensions were confirmed to be produced on the substrate surface, not on the objects, as the imaging was conducted with illumination light under total internal reflection which only excites dyes near to the glass surface. The regions with integrin tension signals are also generally larger than the object sizes. Moreover, the force signals remain on the substrate after the object removal by phagocytosis (Supplementary Video 2). In a control test, RGD-null ITS did not exhibit any fluorescence signal (Extended Data Fig. 4), suggesting the fluorescent signal of ITS is indeed activated by integrin tensions. The existence of integrin tensions suggests that integrin-mediated adhesion structures were formed on the substrates to surround the surface-bound objects during phagocytosis. To our knowledge, this adhesion structure has not been previously reported, and we named it phagocytic adhesion ring (PAR). F-actin was also consistently recruited to PAR. The time-series imaging demonstrates periodic dynamics of F-actin on PAR, while integrin tension signal was steadily accumulated (Fig. 1C and 1D).

## Integrin tensions in PARs contribute to phagocytosis of surface-bound objects

It is clear that integrins in the PAR actively transmit tensions. We ask the question whether the integrin tensions contribute to the phagocytosis of surface-bound objects. First, we recorded the internalization process of surface-bound beads in Supplementary Video 2. The beads were loosely immobilized to allow phagocytosis (See Methods). Integrin tensions were consistently produced around the beads before their internalization, suggesting that integrin tensions likely contribute to the phagocytosis of surface-bound objects. To rigorously demonstrate that integrin tensions facilitate the phagocytosis of surface-bound objects, we developed an assay to quantify the phagocytosis efficiency while tuning the adhesion strength of the beads or integrin ligand strength. Because ITS is irreversible tension sensor that records historic force signal, the force signal in a ring pattern faithfully mark the site of phagocytosis even after the internalization of objects. We imaged PAR force rings and examined the presence of beads on the force rings to confirm whether a bead has been internalized in a PAR. A force ring without a bead is considered as a successful phagocytosis (Fig. 2A). Phagocytosis efficiency is quantified as the percentage of force rings without beads versus all force rings.

**Fig. 2.**
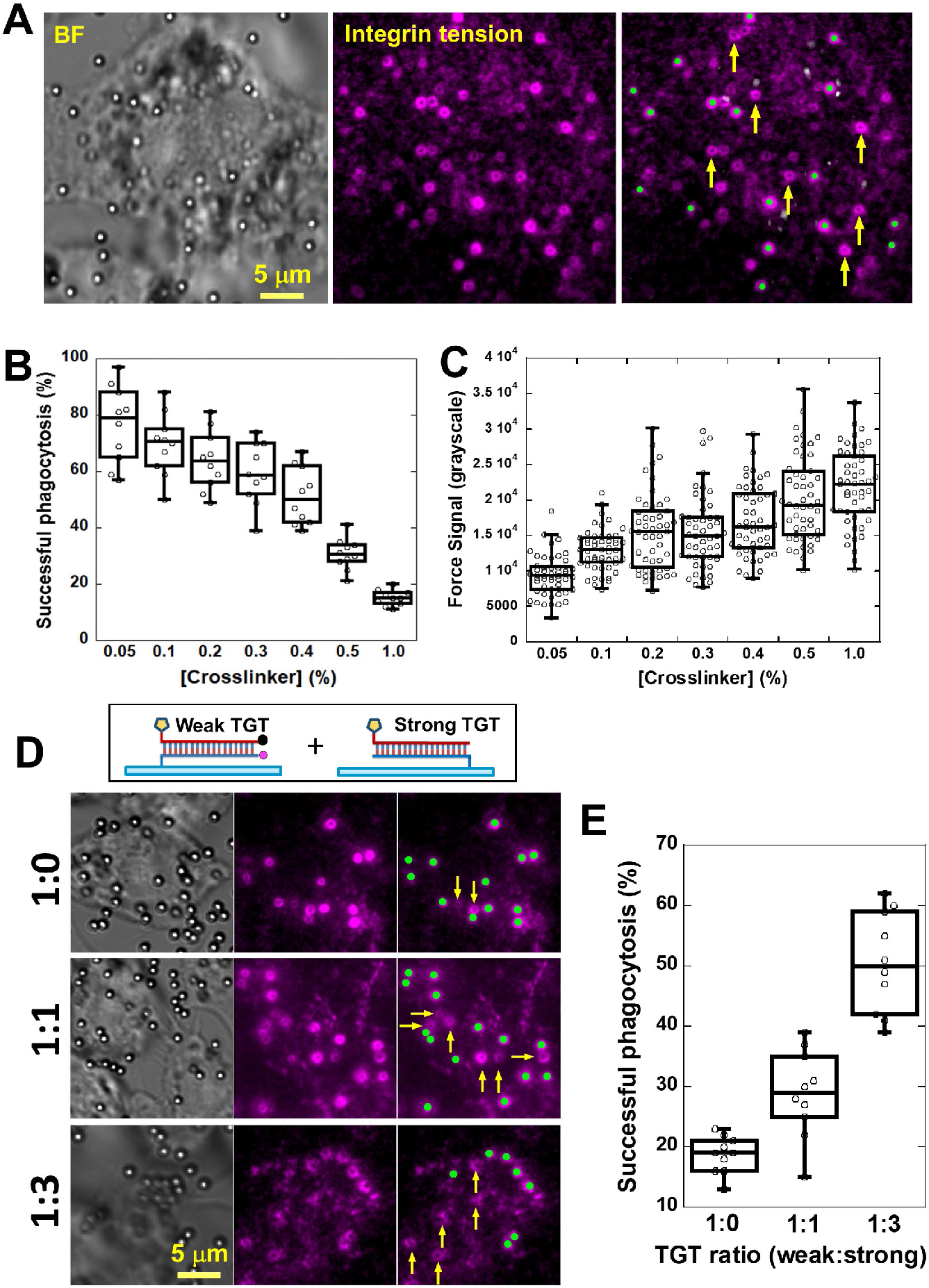
Integrin tensions facilitate the phagocytosis of surface-bound objects. (**A**) A force assay has been developed to quantify phagocytosis efficiency. Green dots mark the presence of beads on force rings. Force rings without beads (Yellow arrows mark a few examples) indicate successful phagocytosis. (**B**) Beads are immobilized on the surface by glutaraldehyde via chemical crosslinking. Phagocytosis efficiencies were quantified on a series of bead-immobilized surfaces. (**C**) Integrin tension signals were quantified on these surfaces. (**D**) TGTs as rupturable integrin ligands, replacing fibronectin, were used to restrict the molecular force level of integrin tensions. Weak TGT (DNA/PNA in unzipping conformation) was mixed with strong TGT (DNA/PNA in shear conformation) at 1:0, 1:1 and 1:3 ratios and coated on the substrates, respectively. Yellow arrows mark the successful phagocytosis. (**E**) Phagocytosis efficiencies versus the ratio between weak TGT and strong TGT.

With this assay, first we tuned the adhesion strength of the beads on the surface. Glutaraldehyde was used to covalently crosslink the beads to the substrate (See Methods). The adhesion strength of beads is tuned by varying the glutaraldehyde concentration. The results showed that phagocytosis efficiency was lower when the bead-substrate crosslinking was stronger (higher glutaraldehyde concentration) (Fig. 2B). Correspondingly, the integrin tension signal intensity became stronger as the bead-substrate crosslinking strength increased (Fig. 2C). This suggests that the intensity of integrin tensions during phagocytosis is correlated with the adhesion strength of the beads.

Next, we sought to investigate the effect of integrin-ligand tethering strength on the phagocytosis efficiency of surface-bound objects. We replaced fibronectin with tension gauge tether (TGT)^18^, a rupturable integrin ligand with tunable tethering strength on the surface. Two TGT constructs in either unzipping or shear conformation were adopted (Fig. D). The critical forces rupturing these two TGTs are expected to be ∼10 picoNewton (pN) and ∼50 pN, respectively^18^. The unzipping-TGT provides weak support for integrin tensions in PARs and restricts integrin tensions under ∼10 pN, whereas the shear-TGT provides strong support for integrin tensions. Both TGTs are synthesized with DNA/PNA duplexes to resist membrane DNase of macrophages. Note that ITS was adopted as the unzipping-TGT in this test, since ITS can function as an unzipping-TGT and also enable integrin tension visualization, which is needed in this assay. The shear-TGT was not labeled with a quencher-dye pair. The two TGTs were mixed at concentration ratios of 1:0, 1:1 and 1:3, respectively, and coated on a surface with beads immobilized by 0.4% glutaraldehyde. Macrophages induced from RAW 264.7 cells were plated on these surfaces. Results show that PARs were formed on all surfaces, suggesting that PAR formation is not sensitive to the force level of integrin tensions sustained by the ligands. However, higher phagocytosis efficiency was observed on the surface grafted with stronger TGT (Fig. 2D-2E). This test demonstrates that integrin tensions in PAR are critical for the phagocytosis of surface-bound objects. Restricting integrin tensions in PAR down to 10 pN (by unzipping-TGT) significantly reduced the phagocytosis efficiency.

## Basic structural proteins in PARs

Next, we investigated the basic protein components in the PAR using immunostaining and fluorescence imaging. Integrin tensions and Rab5, an early phagosome marker^23, 24^, were shown to be co-localized in the PAR (Fig. 3A), suggesting that the PAR is a machinery related to phagocytosis. Integrin β_2_ was confirmed to exist in the PAR (Fig. 3A). Knockdown of integrin β_2_ in the RAW cells by siRNA treatment caused weaker integrin tension signals (Fig. 3B) in the PAR and less efficient phagocytosis (Fig. 3C), suggesting that integrin β_2_ is the major integrin type mediating adhesive force in PAR and subsequent phagocytosis. Talin 1 and vinculin were also recruited to the PAR (Fig. 3A). Because talin and vinculin are well-known force-transmitting proteins in other integrin-mediated adhesion structures such as focal adhesions^25^ and podosomes ^26^, the presence of talin and vinculin in the PAR suggests that integrin tension transmission in the PAR may similar molecular linkage to those in focal adhesions and podosomes.

**Fig. 3.**
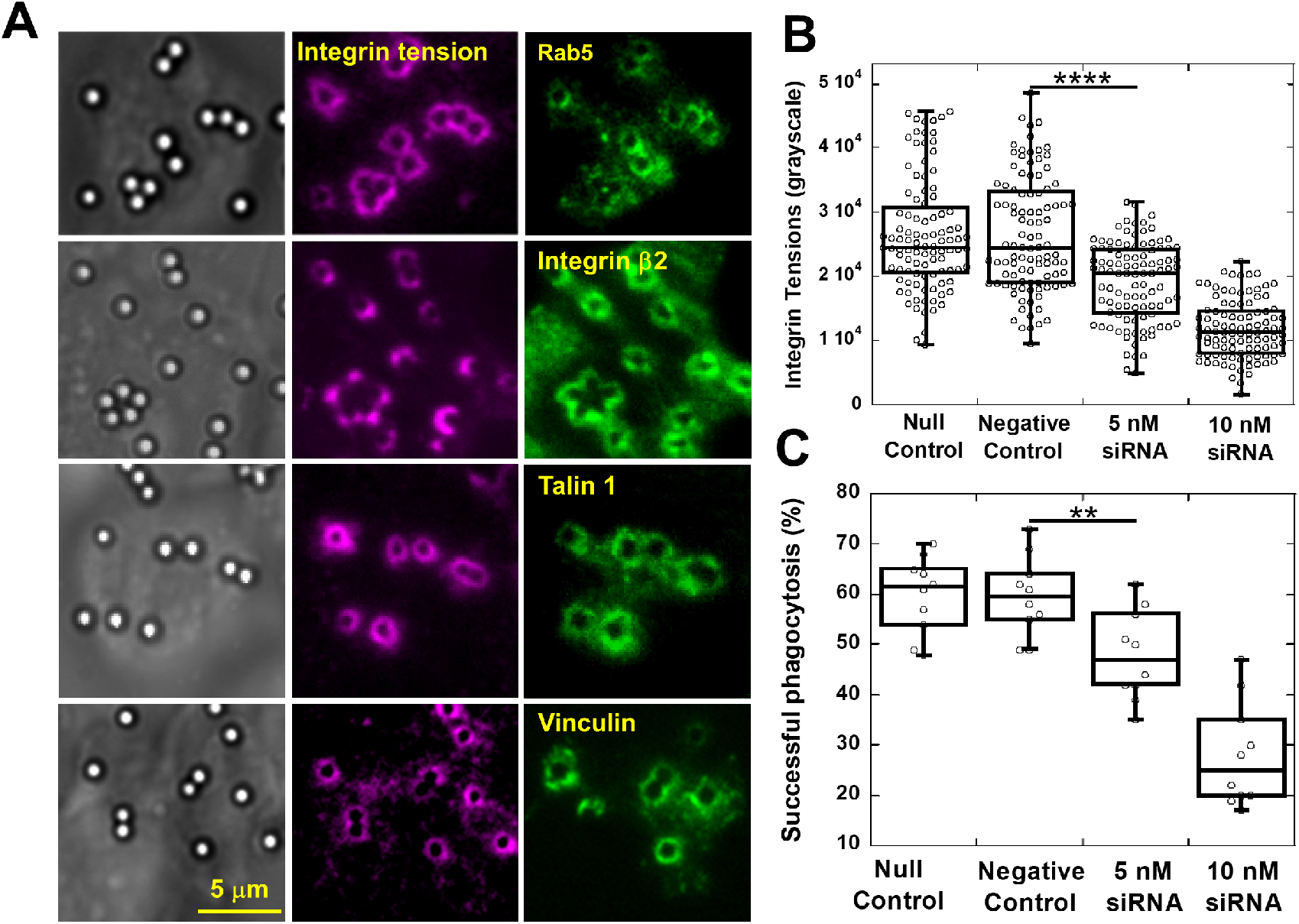
Protein components of phagocytic adhesion rings. **(A)** Rab5, integrin β_2_, talin and vinculin are confirmed in PARs. (**B**) Knockdown of integrin β_2_ by siRNA reduced integrin tension signals in PAR significantly. p value <0.0001 between control group and 5 nM siRNA-treated group. (**C**) Knockdown of integrin β_2_ by siRNA reduced phagocytosis efficiency of surface-bound beads significantly. p value <0.01 between control group and 5 nM siRNA-treated group.

## Integrin tensions in PARs are resulted from F-actin polymerization, but not myosin II

Next, we investigated the force sources of integrin tensions in the PAR. It is known that actomyosin (myosin II coupled actin fibers) is the main source of integrin tensions in focal adhesions ^27^. Here we investigated the role of myosin II in PAR formation and force generation. We incubated macrophages on the microbead-immobilized ITS surface. The macrophages were treated with 25 μM blebbistatin or 20 μM Y27632 which inhibits myosin II function. The results show that the inhibition of myosin II had little effect on the PAR formation or on integrin tension generation in the PARs (Fig. 4A and 4B). In contrast, inhibition of myosin II by blebbistatin abolished FA formation and drastically reduced integrin tension signals in HeLa cells (Extended Data Fig. 5). This result suggests that myosin II does not play a significant role in PAR formation or integrin force generation in PARs. At last, we inhibited actin polymerization with 100 nM Latrunculin A and inhibited Arp2/3 complex (regulating the initiation and branching of actin polymerization) with 100 μM CK666, respectively^28, 29^. The inhibition of actin polymerization or Arp2/3 significantly reduced the force signal in PAR (Fig. 4A and 4B), suggesting that integrin tensions in the PAR were resulted from actin polymerization.

**Fig. 4.**
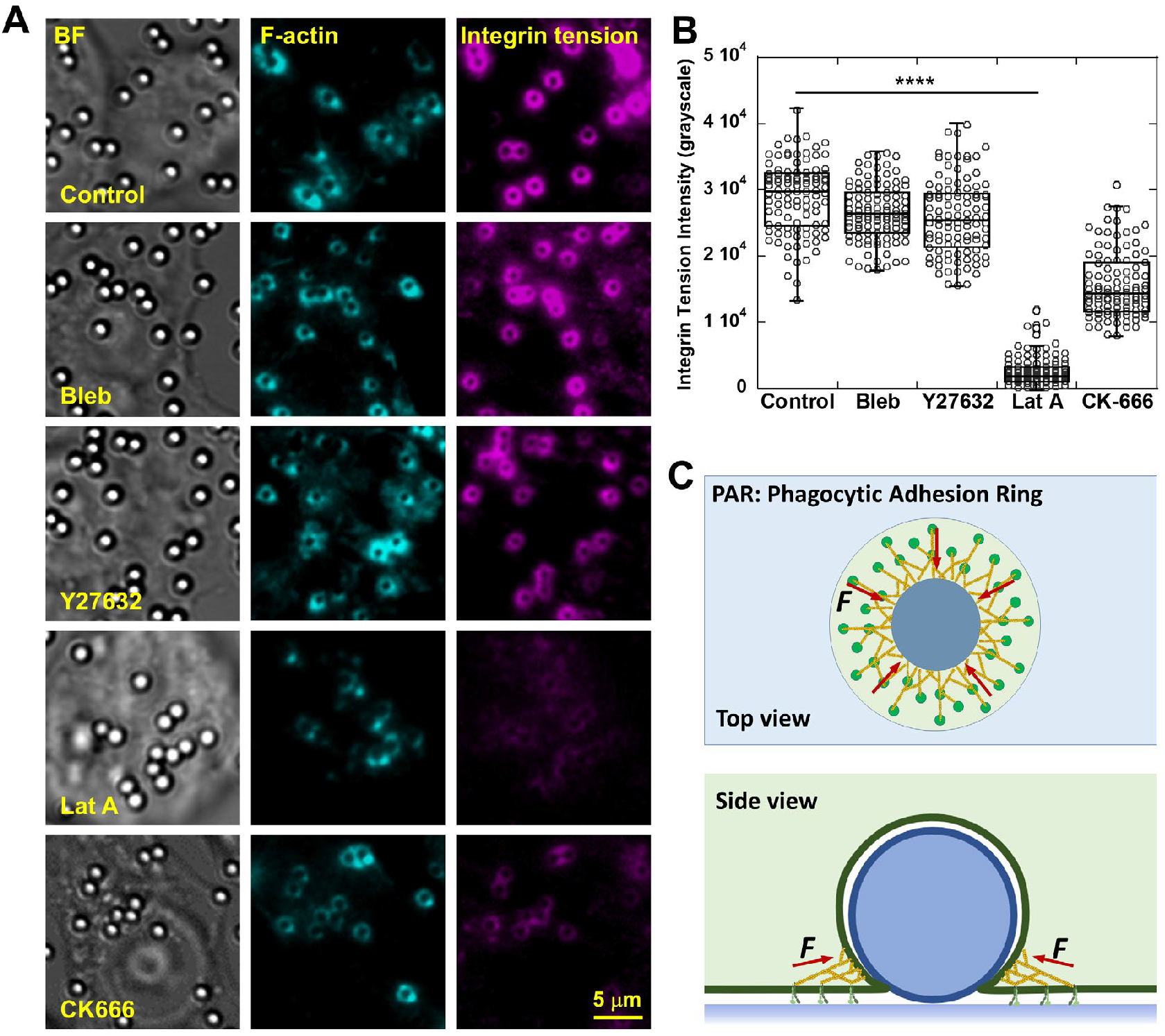
Integrin tensions in PARs originate from actin polymerization. (**A**) Integrin tensions in PARs with the treatment of DMSO (control), 25 μM Blebbistatin (Bleb), 20 μM Y27632, 100 nM Latrunculin A (Lat A) and 100 μM CK666, respectively. (**B**) Intensities of integrin tensions in PARs under different inhibition conditions. 100 PARs were analyzed for each condition. p value < 0.0001 between control group and Lat A-treated group. (**C**) A simplistic model of PAR and PAR-assisted phagocytosis. Actin (yellow) polymerization supported by integrins (green) in PARs squeezes beneath surface-bound objects and lift the objects up. Integrins sustain the reaction force from objects.

## Discussion

Based on above observations, we propose the model of PAR-mediated phagocytosis (Fig. 4C). When macrophages encounter surface-bound objects under the cell bodies, the local actin polymerization experiences obstacles and subsequently recruit integrins to anchorage the local actin network surrounding the objects on the substrate. Integrins continue to recruit other structural proteins and eventually form the PAR structure. Rooted on the PAR, actin polymerizes and squeezes underneath the objects, producing a force that may lift the objects off from the surface,while the counteracting force from the objects is transmitted through actin network to integrins in the PAR, hence producing the integrin tensions. This process does not require myosin II-mediated actin filament contraction. Therefore, inhibition of myosin II has insignificant effect on PAR force generation and phagocytosis of surface-bound objects.

In summary, we revealed a general mechanism of phagocytosis adopted by macrophages to internalize surface-bound objects. As many pathogens can strongly adhere on the tissues of the host, we show that macrophages form unique integrin β_2_-mediated adhesion structures termed PAR on a substrate to encircle surface-bound objects. Integrins in the PARs sustain tensions to support local actin polymerization which squeezes underneath the objects, eventually lifting them off. This mechanism is different from phagocytosis of suspended objects during which macrophages bind to objects in suspension directly by phagocytic receptors^30^, which require the suspended objects to possess corresponding ligands which are either inherently expressed by the pathogens or tagged by the host through opsonization^31, 32^. However, for the internalization of surface-bound objects, direct cell-object binding is not needed and objects are not required to labeled with the ligands for phagocytic receptors. Instead, PAR coupled with F-actin provides a different biomechanical process which, instead of pulling the objects into cells, squeezes into the substrate-object cleavage and pinches the objects off. Considering the fundamental role of macrophages in immunity and pathology, and the ubiquitous surface-bound pathogens and cell debris, we suggest that this new mechanism is important for the understanding of the function of macrophages in immune defense and homeostasis.

## Acknowledgments

This research was supported by National Institute of General Medical Sciences (1R35GM128747). We thank all laboratory members for comments on the study and the manuscript.

## Supplementary Figures

**S. Fig. 1.**
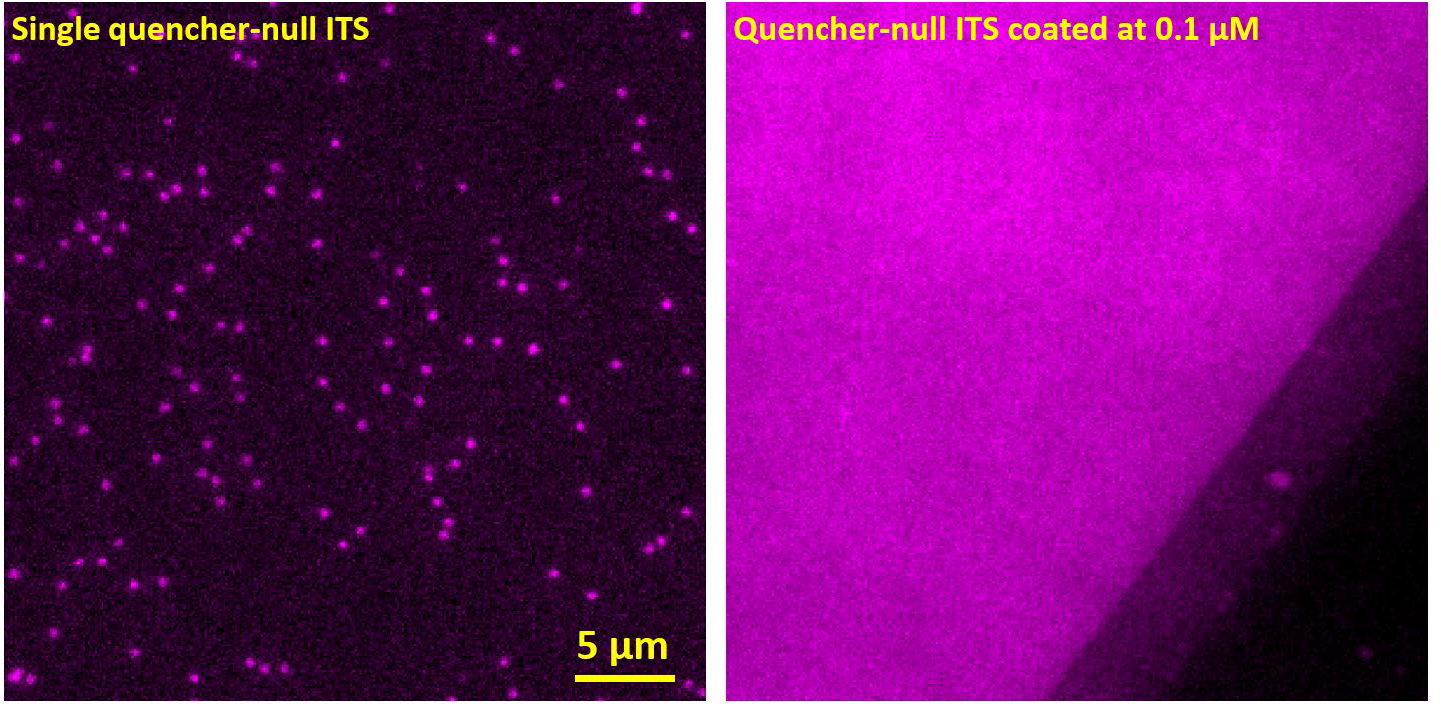
Calibration of ITS surface density. (**A**) Quencher-null ITS (ITS without the quencher) was coated on a pegylated glass surface at a concentration of 5 pM (picomolar). The Cy5 dye in the ITS was imaged at the single molecule level with 1 sec exposure time and 10% laser power. The total fluorescence intensity of 5×5 pixels under single Cy5 dyes was calculated and averaged to be 21456. (**B**) Quencher-null ITS coated at 0.1 μM (concentration used in the phagocytosis assay) was imaged with 10 ms exposure time and 10% laser power. Fluorescence intensity of single pixel (corresponding to 0.11×0.11 μm^2^ sample area) was calculated and averaged to be 3284. ITS molecular density on the glass surfaces was therefore evaluated as 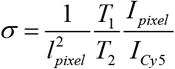, where *l* _*pixel*_ is 0.11 μm of a pixel, *T*_1_ is exposure time of single molecule imagine (1 sec), *T*_2_ is exposure time of non-single molecule imagine (10 ms), *I*_*Cy*5_ is the total fluorescence intensity of single Cy5s and *I* _*pixel*_ is the fluorescence intensity on a single pixel. The coating density of ITS was evaluated to be 1265/μm^2^ for the phagocytosis assays.

**S. Fig. 2.**
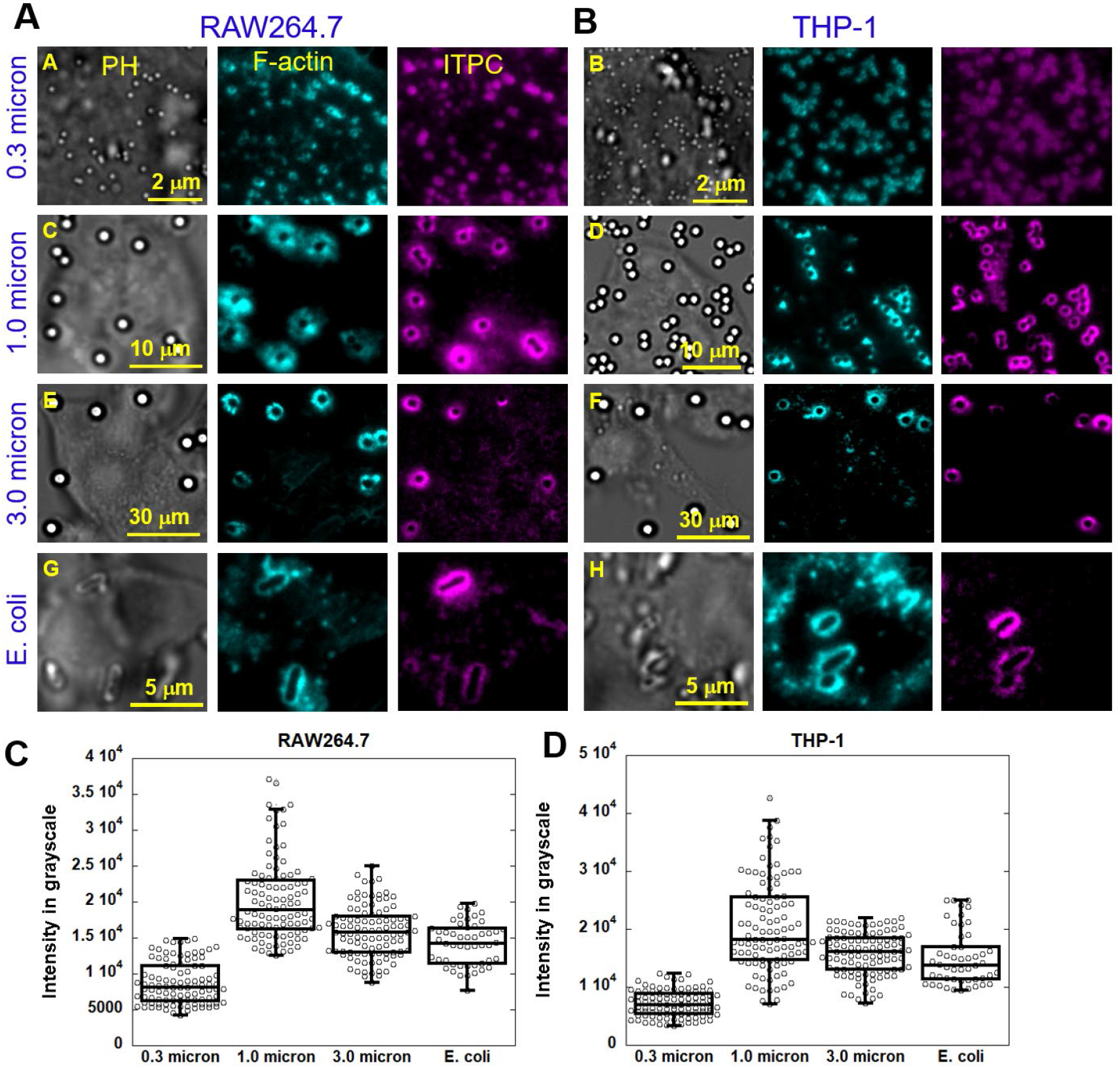
Co-imaging of integrin tensions and F-actin in RAW and THP-1 cells on particles with various sizes. **(A)** RAW and **(B)** THP-1 macrophages were incubated on ITS modified surface against microbeads with diameter of 0.3 μm, 1 μm and 3 μm and E. coli respectively. F-actin was stained with dye-labeled phalloidin. (**C-D**) Quantification of integrin tension signal intensities against the surface-bound objects. Both cells have shown integrin tensions around the objects regardless of the sizes and nature of the targets.

**S. Fig. 3.**
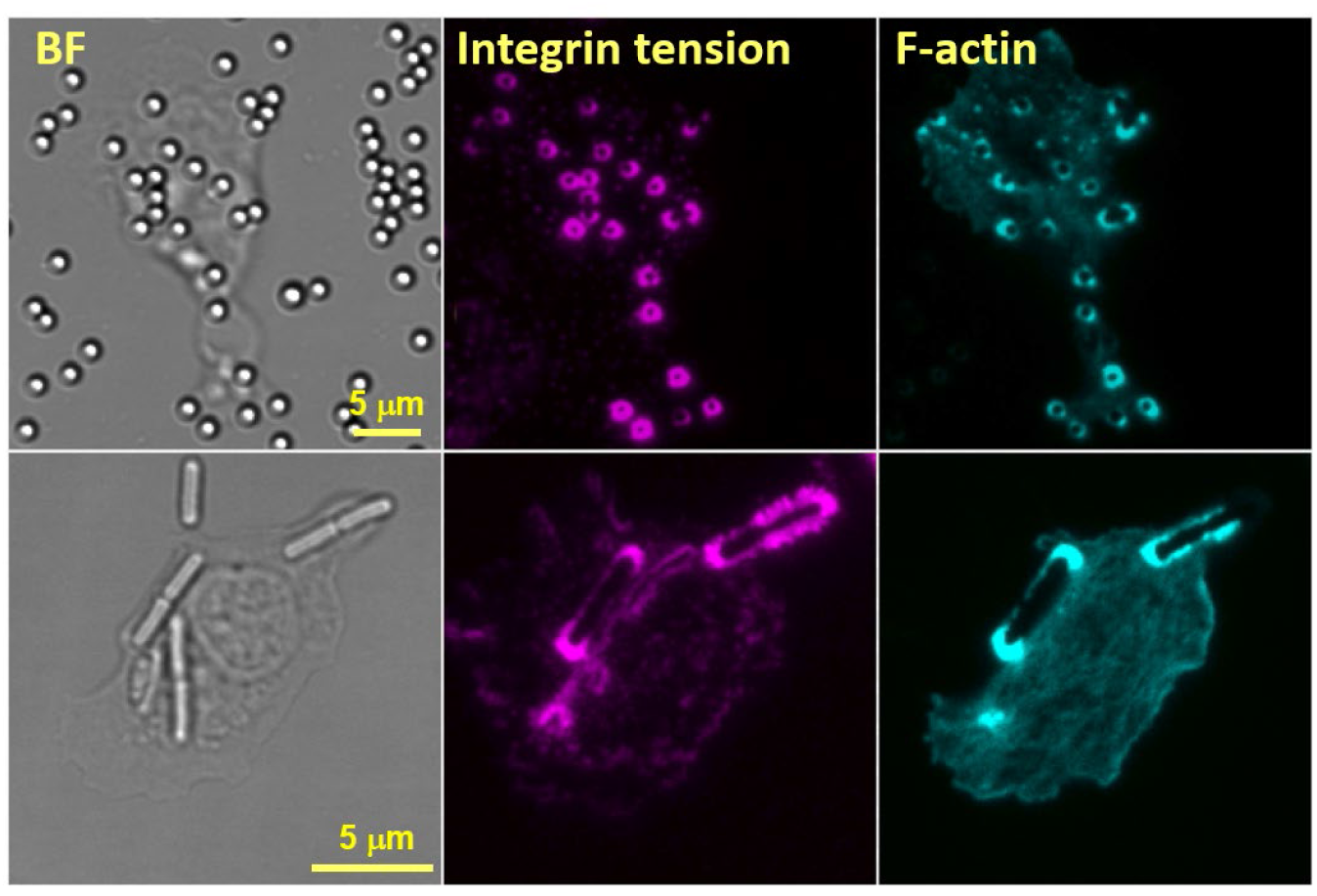
Macrophages extracted from fish exhibited PAR formation and produced integrin tensions in the PAR.

**S. Fig. 4.**
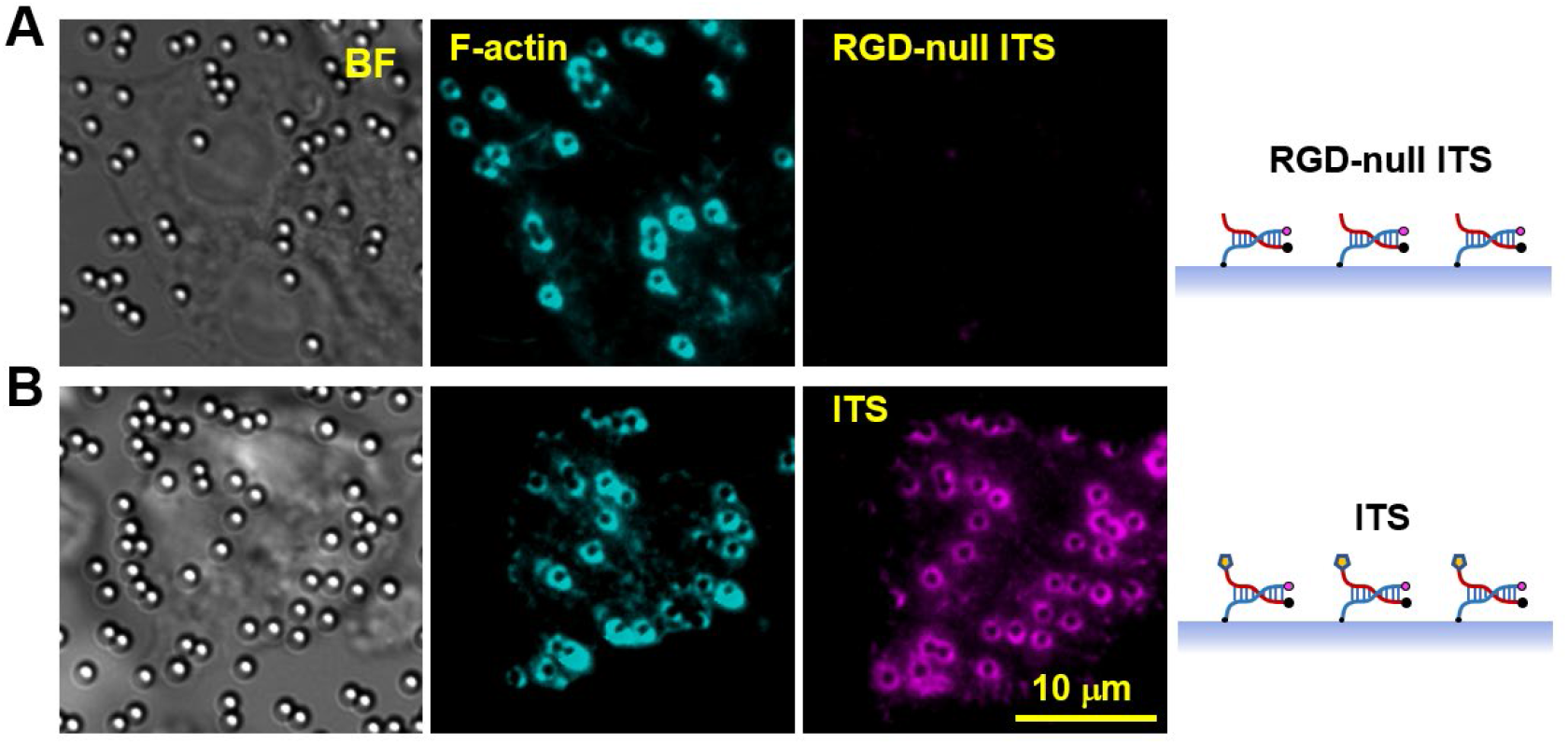
ITS reports integrin-transmitted tensions. **(A)** An ITS construct without RGD (integrin ligand) did not show any fluorescence signal despite the clear formation of PAR indicated by F-actin. Note the surface is also coated with fibronectin to support PAR cell adhesion and PAR formation. **(B)**. ITS with RGD reports ring-shaped force patterns.

**S. Fig. 5.**
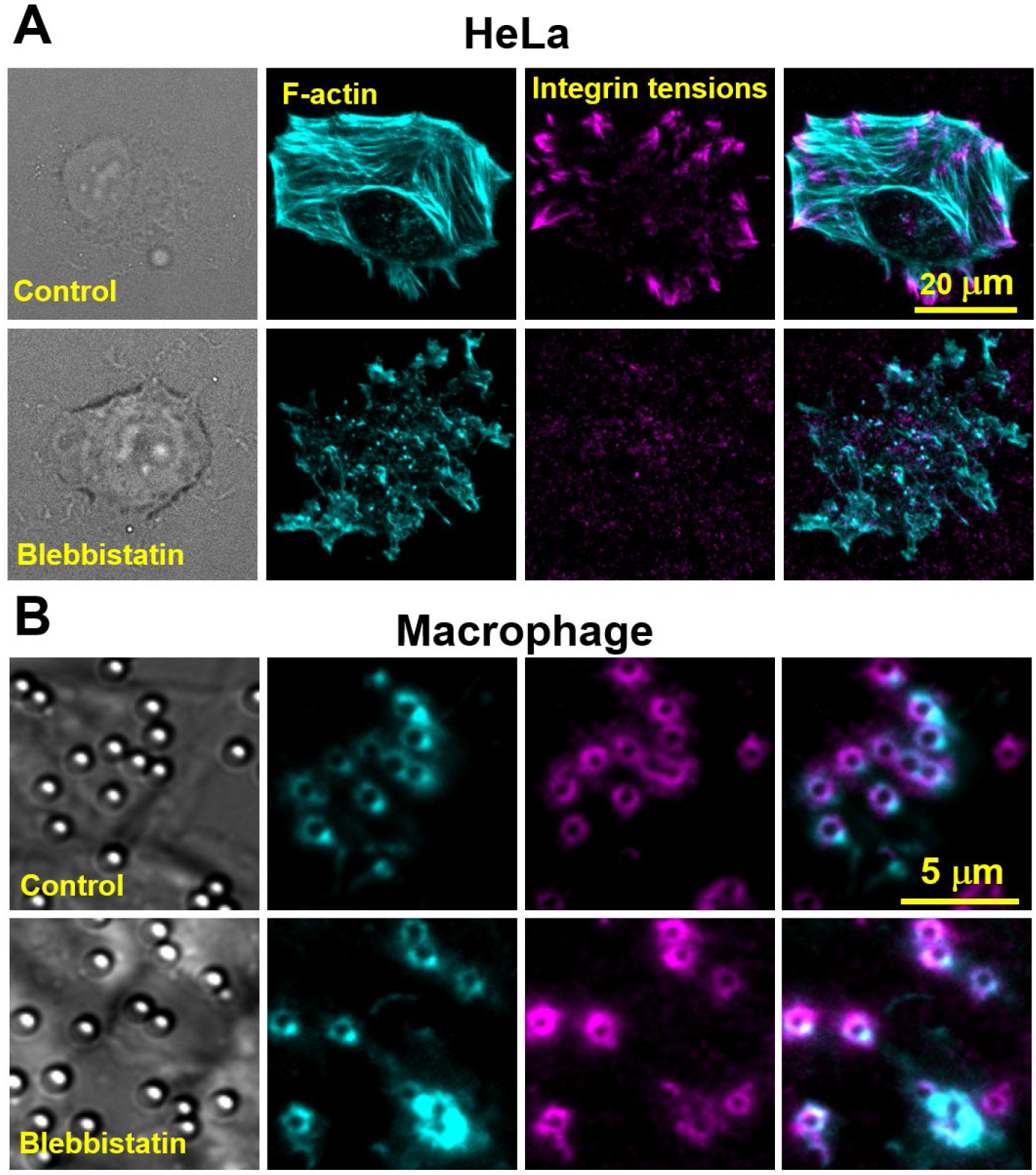
Contrasting effect of myosin II inhibition on integrin-tensions in focal adhesions (FAs) and in PAR. (**A**) Co-imaging of F-actin and integrin tensions in HeLa cells (forming FAs) treated with DMSO (control) or 25 μM blebbistatin, respectively. (**B**) Co-imaging of macrophages (RAW 264.7) treated with DMSO (control) or 25 μM blebbistatin, respectively. Myosin II inhibition abolished FA formation and reduced integrin tensions in FAs, but it had insignificant impact to PAR formation or integrin tension generation in PARs.

